# A small molecule increases mycobacterial translation fidelity without a fitness cost by targeting ribosomal protein S5

**DOI:** 10.1101/2024.10.20.619312

**Authors:** Jie Wu, Swarnava Chaudhuri, Sarah Caldwell Feid, Hemant Joshi, Miaomiao Pan, Qingwen Kawaji, Gang Liu, Babak Javid

**Affiliations:** Tsinghua University; UCSF

**Keywords:** mistranslation, translational fidelity, small molecule, antibiotic tolerance, medicinal chemistry, mycobacteria

## Abstract

A central dogma of molecular biology is the "speed-accuracy trade-off," where ribosomes must slow down to ensure accurate protein synthesis. In mycobacteria, a high basal level of mistranslation at glutamine and asparagine codons, caused by an indirect tRNA aminoacylation pathway, promotes tolerance to the antibiotic rifampicin. While pharmacologically increasing translational fidelity is a promising strategy to combat antibiotic tolerance, the underlying mechanisms remain poorly understood. Here, we screened 9,000 synthetic compounds and identified **benzo[d]isoxazole-4,7-diones** as a novel chemical class that reduces mycobacterial mistranslation. Medicinal chemistry optimization yielded a lead compound, **9787**, with superior potency in decreasing mistranslation and reversing rifampicin tolerance. Using competitive chemical proteomics, we identified the 30S ribosomal protein S5 (RpS5) as the specific cellular target. Remarkably, compound **9787** enhances translational fidelity at concentrations that do not measurably impact the overall rate of protein synthesis. Our findings challenge the universality of the speed-accuracy trade-off, demonstrating that fidelity can be improved independently of translation speed. This work reveals that the ribosomal small subunit is a druggable target for modulating translational quality control and introduces a new strategy for combating antibiotic-tolerant bacteria without the associated fitness cost of slowed translation.

**Importance:** A fundamental principle in molecular biology holds that ribosomes face a trade-off between translation speed and accuracy: going faster means making more errors, while maintaining high fidelity requires slowing down. This study challenges that paradigm by identifying a small molecule that increases translational accuracy in mycobacteria without affecting the rate of protein synthesis. The compound targets ribosomal protein S5 and specifically reduce errors arising from physiologically mischarged tRNAs – a quality control problem distinct from the well-studied codon•anticodon mismatches. This form of mistranslation contributes to antibiotic tolerance in tuberculosis, making it a potential therapeutic target. Our findings reveal unexpected flexibility in how ribosomes maintain translation quality and suggest that pharmacologically increasing fidelity without the fitness cost of slowed protein synthesis may be an attractive strategy for combating antibiotic-tolerant bacteria.

## INTRODUCTION

Genetically susceptible bacteria may still withstand killing by antibiotics, a phenomenon known as antibiotic tolerance (1, 2). There are several potential mechanisms for tolerance, and these may reflect different physiological states of the bacteria (3-5). Non-replicating persisters are usually multi-drug tolerant and have a relatively, but not entirely, quiescent physiological state (5-8). We recently described a form of tolerance in mycobacteria to the first-line antibiotic rifampicin, which we termed RNA polymerase-specific phenotypic resistance (RSPR) (9-11). By contrast with non-replicating persisters, the antibiotic-tolerant bacteria in RSPR are able to grow in bulk-lethal concentrations of antibiotic that can kill the majority of bacilli in the culture (9). Importantly, RSPR was apparent in *Mycobacterium tuberculosis* freshly isolated from sputum from a small number of patients, suggesting that it may be a clinically relevant mode of antibiotic tolerance (10, 12).

We had identified at least two mechanisms that contribute to mycobacterial RSPR (10, 11). One of these mechanisms involved modulation of translational fidelity in the indirect tRNA aminoacylation pathway – or specific mycobacterial mistranslation (10, 13). Unlike cytosolic eukaryotic gene translation, the vast majority of bacteria, excepting *Escherichia coli* and a few related proteobacteria, lack the specific tRNA aminoacyl synthetases for either glutamine or asparagine or both (14-18). All mycobacteria lack both glutaminyl- and asparaginyl-synthetases. Instead, they undergo a two-step pathway to specifically aminoacylate glutamine and asparagine tRNAs. First, a non-discriminatory glutamyl-(or aspartyl-) synthetase physiologically misacylates the glutamine or asparagine tRNA respectively to Glu-tRNA^Gln^ and Asp-tRNA^Asn^. This misacylated complex is then specifically recognized by the essential glutamine amidotransferase GatCAB (10). Using cytosolic glutamine as an amide donor, GatCAB amidates the misacylated tRNAs to their cognate Gln-tRNA^Gln^ and Asn-tRNA^Asn^ respectively, thereby allowing preservation of the genetic code. Therefore, mutations that alter GatCAB structure and stability can increase mistranslation in mycobacteria (10, 13). Moreover, we had identified that naturally occurring mutations in *gatA* in clinical isolates of *Mycobacterium tuberculosis* both increased rates of specific mistranslation and tolerance to rifampicin (10). Furthermore, GatCAB could be limiting in wild-type mycobacteria, and sorting cells by decreased expression of wild-type *gatCAB* in an isogenic population identified subpopulations with both increased rates of mistranslation and antibiotic tolerance (10). We also showed that mistranslation of a specific, critical, residue in the rifampicin binding region of the cellular target of rifampicin, RNA polymerase (RNAP), contributed substantially to RSPR in wild-type mycobacteria (10). Together, these data compellingly suggested that specific mycobacterial mistranslation was an important contributor to RSPR.

Since mycobacterial mistranslation contributed to RSPR even in isogenic wild-type bacteria, we hypothesized that this pathway may be amenable to pharmacological intervention. We identified the natural product kasugamycin as a small molecule that could both decrease specific mycobacterial mistranslation and potentiate rifampicin both *in vitro* and in a murine model of tuberculosis (TB) (19). However, other than the naturally occurring peptide Edeine, which showed some activity in our assays (19), no other molecules capable of decreasing this specific form of bacterial mistranslation, which is mechanistically distinct from ribosomal decoding errors (10), have been identified. We decided to perform a pathway-specific whole-cell screen (20) that could identify synthetic small molecules that could target mycobacterial mistranslation. In this study, we utilized our mistranslation inhibition assay, which had earlier been used to identify kasugamycin’s ability to increase mycobacterial translational fidelity (19), (21), in a high-throughput format. Here, we identify analogues of benzo[d]isoxazole-4,7-diones that can decrease mistranslation rates and potentiate rifampicin *in vitro*. Furthermore, we identify the potential target of the most potent analogue as the small ribosomal subunit protein RpS5. Importantly, the lead compound, **9787**, increases translational fidelity at concentrations that do not impact the rate of protein synthesis, challenging the universality of the speed-accuracy trade-off and revealing previously unrecognized mechanisms of ribosomal quality control.

## RESULTS

### Development of a mistranslation pathway-specific phenotypic screen

We developed a reporter system in *M. smegmatis* to sensitively measure relative mistranslation of asparagine to aspartate – one of the two form of mistranslation implicated in the indirect tRNA aminoacylation pathway (**Fig. 1A**) (19), (21). This whole cell reporter system avoids a common pitfall of *in vitro* screens: such screens identify many hits that are ultimately inactive against whole cells due to the complex and thick mycobacterial cell wall and envelope (20). However, unlike a traditional whole-cell screen in which all hits that could kill bacteria are identified, leading to ‘re-discovery’ of common anti-microbials or identification of non-specifically toxic molecules, the system retains the advantage of being “pathway specific”. Only hits that decrease specific mycobacterial mistranslation are identified.

**Figure 1.**
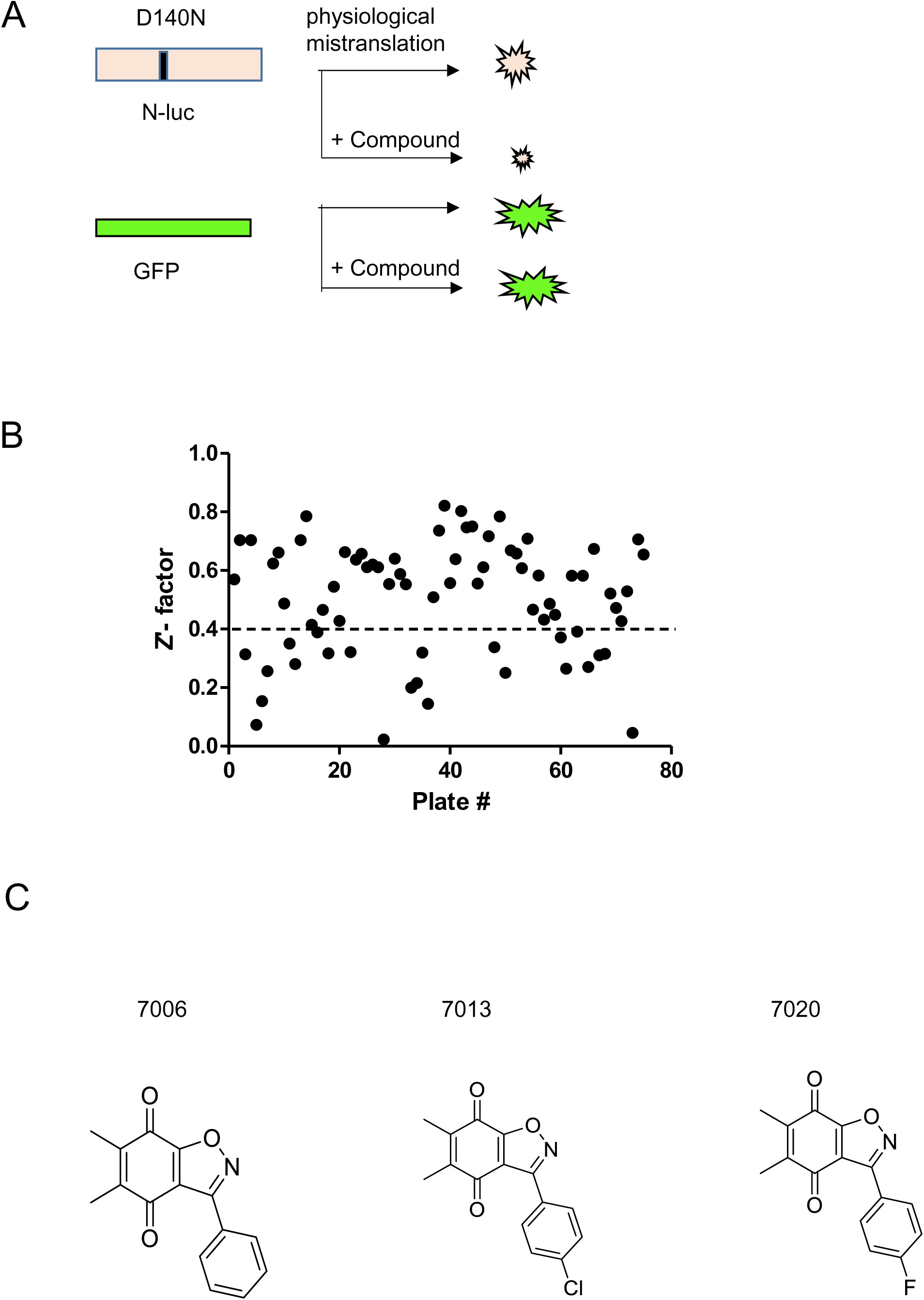
A phenotypic whole-cell screen identified benzo[d]isoxazole-4,7-dione analogs as modulators of mycobacterial mistranslation. (A) Schematic of the N-luc luciferase/GFP reporter system for screening of chemicals that decrease mistranslation in *Mycobacterium smegmatis*. (B) Screening of chemical library using the N-luc luciferase/GFP reporter for inhibitors of mycobacterial mistranslation. (Materials & Methods for details) The quality of the assay was assessed by the Z-factor, which was calculated by the formula 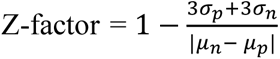, where σ_p_, σ_n_, μ_p_ and μ_n_ are standard deviations (σ) and means (μ) of the positive (p) and negative (n) controls respectively. Throughout the screening process, kasugamycin (Ksg) and dimethyl sulfoxide (DMSO) have been used as the positive and negative controls respectively. (C) Structure of benzo[d]isoxazole-4,7-dione analogs identified in the preliminary screen (7006, 7013, 7020).

The basis of the screen was as follows: Nluc luciferase was mutated at a critical aspartate residue (D140) to asparagine, that resulted in >99% loss of activity (19, 21). Physiological mycobacterial mistranslation of asparagine->aspartate resulted in a baseline detectable Nluc activity of ∼1% of the wild-type enzyme (corrected for cell density and protein expression by normalizing to GFP fluorescence). If a small molecule *decreased* mistranslation rates, the corrected Nluc/GFP would be reduced below the baseline physiological rates (**Fig. 1A**). Hits were classified as those molecules that decreased mistranslation rates by > 1 standard deviation compared with the negative control.

We then used this strain to screen 9000 synthetic heterocyclic small molecules in 96 well-plate format (**Fig. 1B**). Screen performance was variable, due to the inherent noisy signal in measuring low rates of translational error, with 30% of plates yielding Z’ < 0.34 and excluded from analysis. From well-performing plates, we identified 45 hits. Of note, 3 were from the same chemical scaffold, benzo[d]isoxazole-4,7-diones (**Fig. 1C**). These 3 compounds, **7006**, **7013** and **7020**, were retested for their ability to reduce mycobacterial mistranslation and RSPR, confirming them as valid hits (**Fig, 2A and 2B**).

### Medicinal chemistry identifies potent analogs that increase specific translational fidelity

We prioritized **7020** for medicinal chemistry because it modulated translational fidelity without direct antimicrobial activity and was amenable to synthetic modification. This would allow us to deconvolute antimicrobial activity from anti-mistranslation activity. We synthesized 110 analogs of **7020**, of which 10 compounds (**Fig. 3**), showed superior potency to the parent compound, **7020**, both in terms of decreasing mistranslation, and potentiating rifampicin in an assay used to measure RSPR (**Table 1**). The compounds also had minimal direct antimicrobial activity by themselves (**Table 1**), making them suitable for dissecting the contribution of targeting mycobacterial mistranslation on antibiotic tolerance without confounding issues due to inhibitory activity.

**Figure 2.**
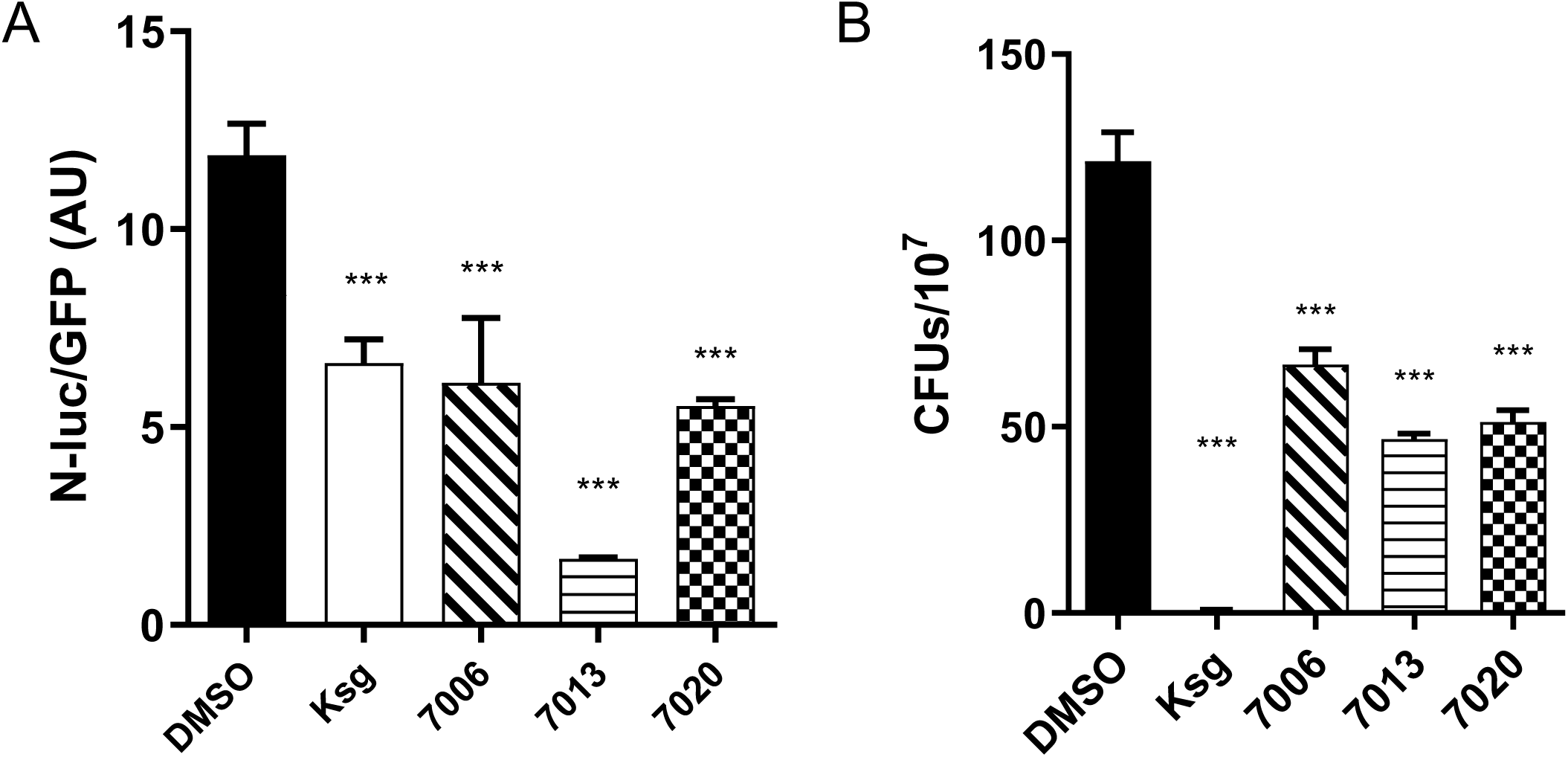
The benzo[d]isoxazole-4,7-dione analogs 7006, 7013 and 7020 decrease mycobacterial mistranslation and RSPR. (A) The compounds 7006, 7013 and 7020 could significantly decrease mistranslation in *M.smegmatis* expressing the N-luc/GFP reporter system. (B) *M. smegmatis* was plated on rifampicin containing media in presence 7006/ 7013/ 7020. All of them could significantly enhance rifampicin mediated mycobacterial killing. DMSO and Kasugamycin (Ksg) have been used as negative and positive controls in both the experiments. ***p < 0.001 by Student’s t-test.

**Figure 3.**
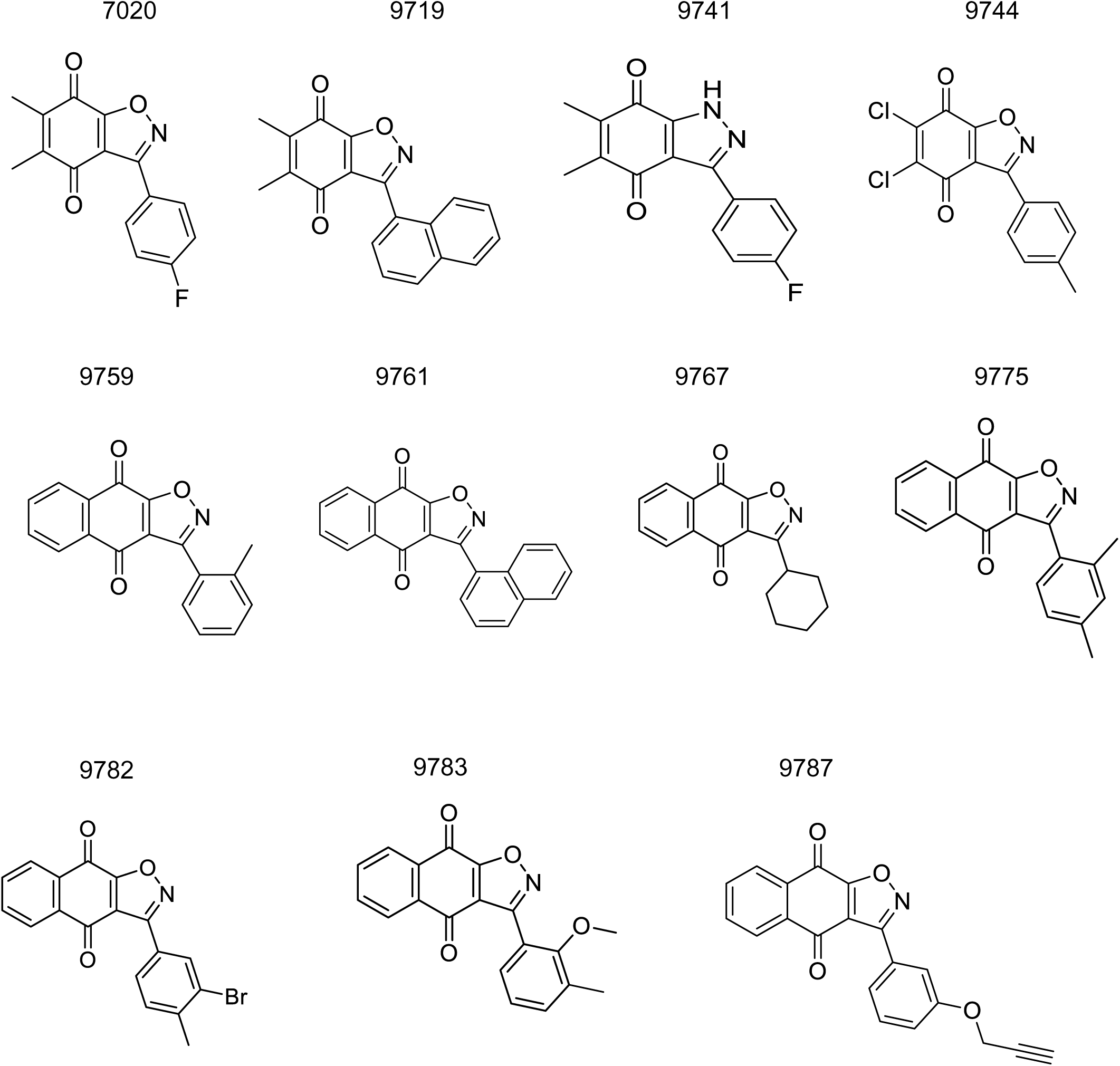
Structural analogs of 7020 which decrease mistranslation and RSPR in *M. smegmatis.* Several analogs of 7020 were synthesized, of which 10 compounds were found to be better than 7020 in inhibiting mycobacterial mistranslation and RSPR (Table 1). The structures of the analog are shown here along with the parent compound 7020.

**Table 1.** Bioactivity of analogs of 7020 with superior potency.

| Compound | Mistranslation Inhibition (%) | RSPR Inhibition (%) | Bacterial Growth Restriction in the Absence of Rifampicin (%) |
| --- | --- | --- | --- |
| 7020 | 53.24 | 57.69 | 0 |
| 9719 | 83.21 | 61.23 | 0 |
| 9741 | 74.38 | 78.34 | 0 |
| 9744 | 86.95 | 87.32 | 1 |
| 9759 | 59.64 | 86.02 | 0 |
| 9767 | 64.77 | 81.43 | 1.2 |
| 9775 | 86.89 | 71.35 | 0 |
| 9782 | 83.73 | 86.78 | 0 |
| 9783 | 66.99 | 88.93 | 0 |
| 9787 | 75.51 | 98.14 | 1.15 |

Our preliminary investigation of structure-activity relationships (SAR) is summarized in **Fig. 4** and **Fig. 5**. Our SAR analysis showed that the benzoquinone part of the compound is essential for its ability to reduce mistranslation; all analogs of 7020 synthesized without the benzoquinone structure are poor inhibitors of mistranslation (**Fig. 4**). We further observed that the methyl groups at positions 5 and 6 of the benzo[d]isoxazole-4,7-diones scaffold are not necessary for their bioactivities in terms of inhibition of mistranslation and RSPR (**Fig. 5**). Replacement of these methyl groups with other functional groups such as chlorine, as in **9744**, or with an aromatic ring, as in **9775**, **9782**, **9783** and **9787**, improved bioactivities of the compounds. (**Table. 1** and **Fig. 5**).

**Figure 4.**
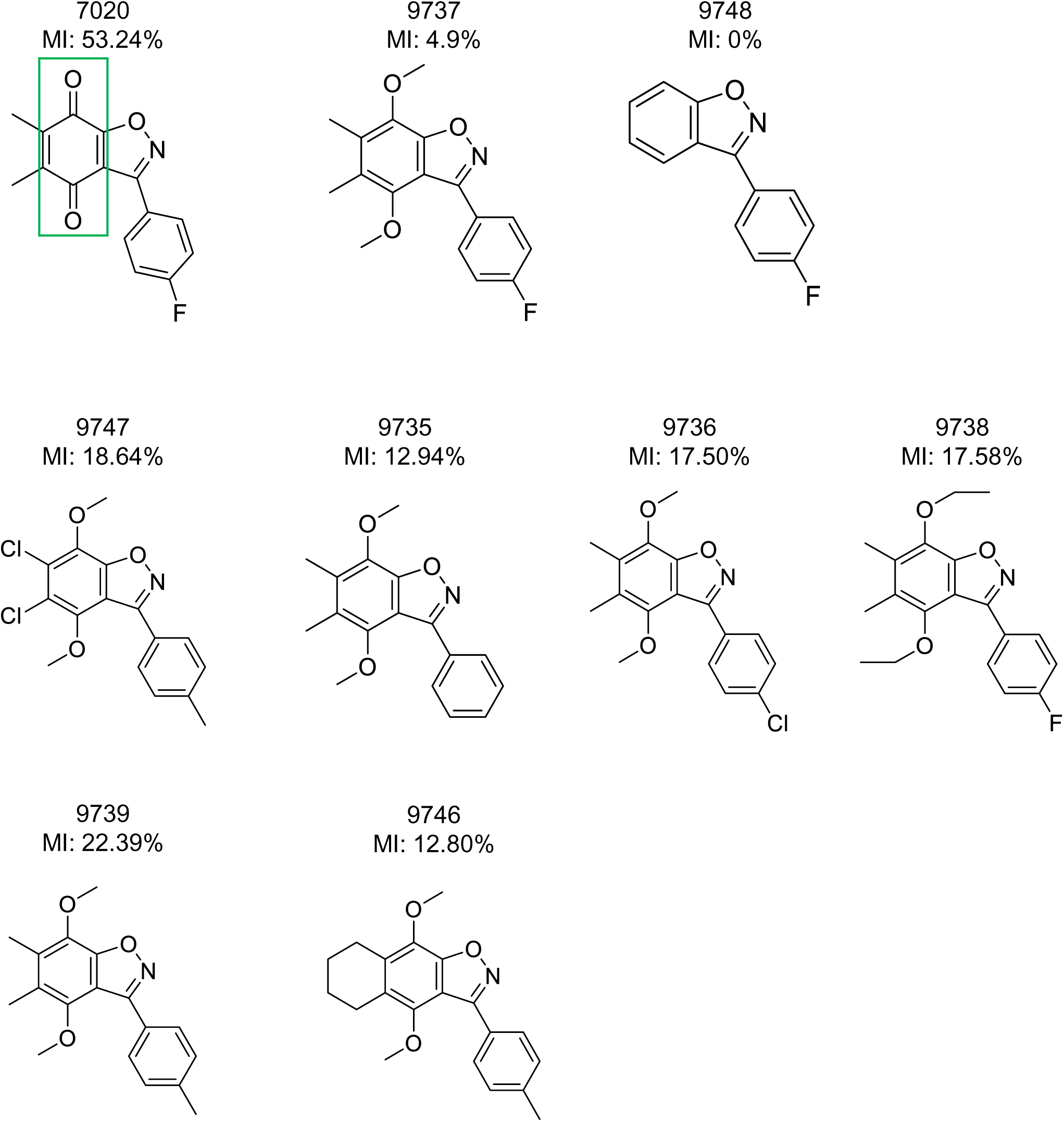
Analogs of 7020 lacking the benzoquione structure are poor inhibitors of mistranslation. The structure of analogs of 7020 synthesized without the benzoquinone structure (boxed in green on 7020) and their percent inhibition of mistranslation are shown alongside 7020.

**Figure 5.**
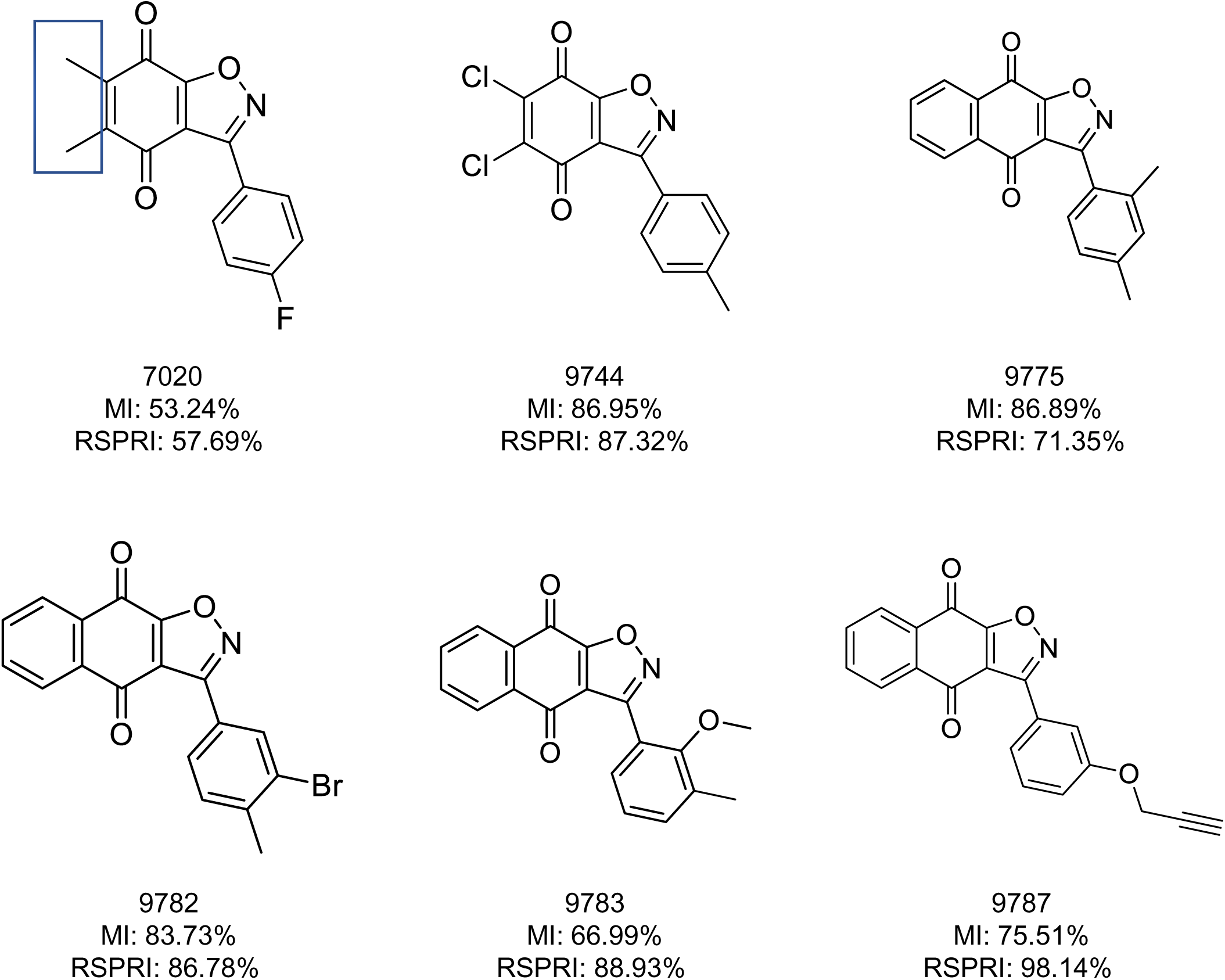
Replacement of 7020 methyl groups improves bioactivity of analog compounds. The structures and percent inhibition of mistranslation and RSPR inhibition of 7020 analogs with improved bioactivity that have replacements of the methyl groups at positions 5 and 6 (boxed in blue on 7020) are shown alongside 7020. In 9744, the methyl groups have been replaced by chlorine. In 9775, 9782, 9783 and 9787, the methyl groups have been replaced by an aromatic ring.

We chose **9787** as the most promising candidate for attempting to determine the cellular target(s) of the hits. **9787** was more potent than the initial hit **7020** (**Fig. 6A**). Moreover, **9787** was able to decrease mistranslation in another closely related mycobacterial species, *M. bovis*-BCG (**Fig. 6B**), demonstrating activity across mycobacterial species.

**Figure 6.**
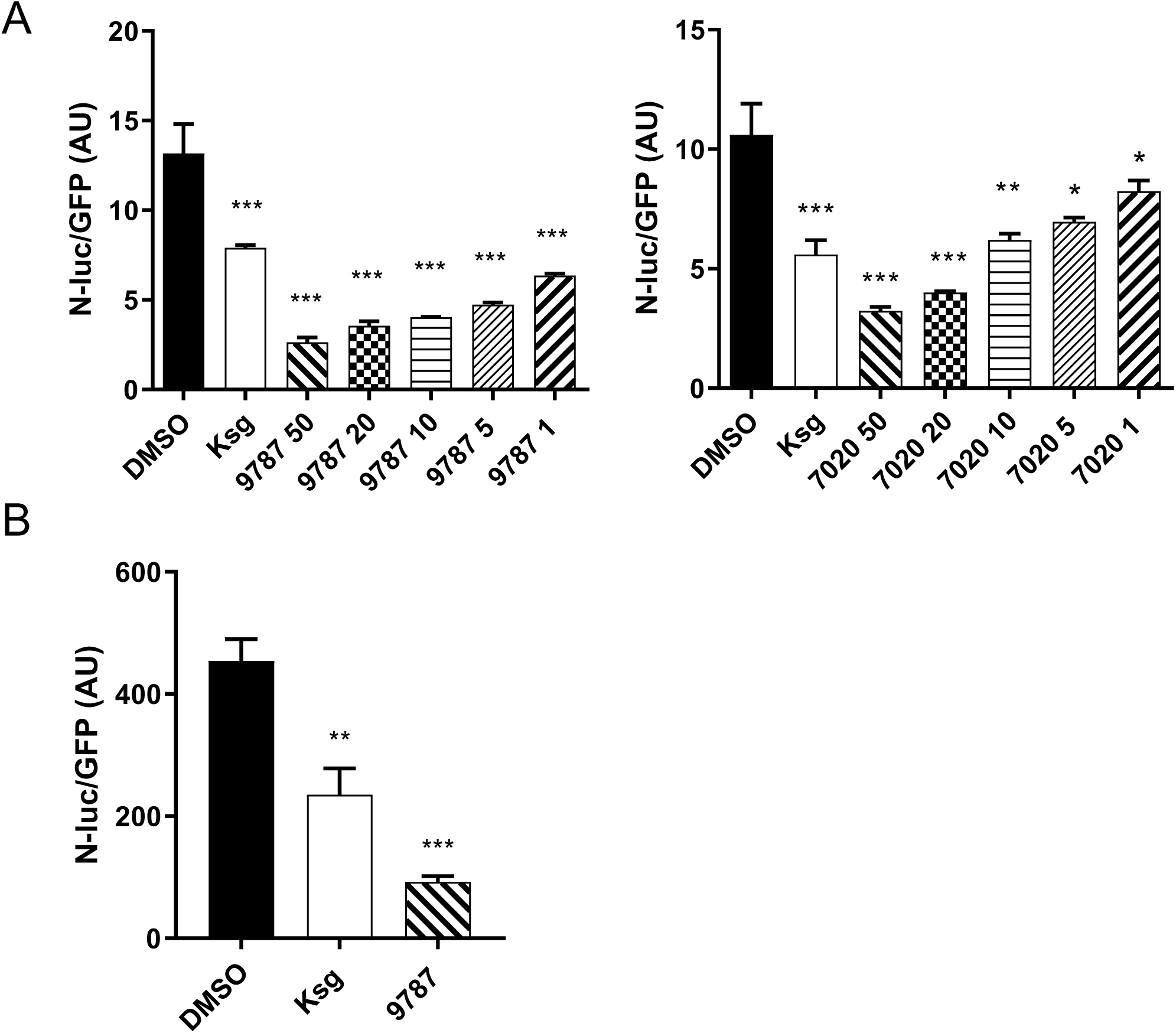
Compound 9787 decreases mistranslation more effectively than compound 7020 and decreases mistranslation in *M. bovis* BCG. (A) The N-luc/GFP reporter was used to show the ability of 9787 to inhibit mycobacterial mistranslation. Different doses of 9787 and the parent compound 7020 were used to check the improved potency of 9787. Dimethyl sulfoxide (DMSO) and kasugamycin (Ksg) have been used as the negative and the positive controls respectively. *p < 0.05, **p < 0.01, ***p < 0.001 by Student’s t-test (B) Other than *M. smegmatis*, the N-luc/GFP mistranslation reporter could be expressed in *M.bovis* BCG strain. 9787 was able to decrease mistranslation in this strain as well. **p < 0.01, ***p < 0.001 by Student’s t-test.

### Chemical proteomics identifies ribosomal subunit protein S5 as the molecular target of 9787

We wished to identify the cellular target by which **9787** mediated a decrease in specific mistranslation rates. Since the compounds lacked significant antimicrobial activity by themselves, it was not possible to raise mutants resistant to the compound and sequence for mutations in the genome to identify potential targets (22). We therefore decided to identify the target by a modified pull-down and mass-spectrometry technique (22-25). For this competitive chemical proteomic approach, **9787** was biotinylated to yield **9788**, which retained activity (**Fig. 7A, B**). An inactive analog, **9789** from the initial medicinal chemistry approach was also biotinylated to make inactive **9791** (**Fig. 7A, C**). We incubated **9788** or **9791** with mycobacterial cell lysate. To increase the specificity of the identified target protein(s), we also included conditions in which 5-fold molar excess of non-biotinylated **9787** or **9789** were added to the tubes in addition to the biotinylated compound (**Fig. 7D** and see methods). Following incubation, the biotinylated compounds and putative cellular binding partners were pulled down using a streptavidin column, washed, and then the eluates were resolved on an SDS-PAGE gel and bands cut out and sent for analysis by mass-spectrometry (**Supplementary Data 1**). Our rationale for identifying the target for **9787** was as follows: When the active biotinylated compound (**9788**) was used, the quality score of the target protein should be significantly higher than that obtained with the inactive biotinylated compound (**9791**). Moreover, in the presence of competing non-biotinylated compounds, less target protein(s) are expected to bind to the streptavidin column. Hence the quality score of the target protein obtained with the active **9788** alone should be significantly higher than that obtained with a mixture of **9788** and excess **9787**. Furthermore, no such phenomenon should be observed in case of the inactive compound (**9789**) and its biotinylated form (**9791**). We plotted the ratio of protein quality scores obtained with the active and inactive biotinylated compounds (**9788/9791**) against the ratio of protein quality scores obtained with **9788** alone, and **9788** and excess **9787** (9788/[9788 + excess 9787]). Out of 1925 proteins, only the 30S ribosomal protein S5 (Rps5) derived from *rpsE* gene, could fulfill all the criteria and significantly stood out against the others. (**Fig. 7E**). These data strongly suggest that RpS5 is the target of active **9787**.

### Compound 9787 increases translational fidelity without affecting translation rate

A fundamental dogma in translation biology holds that increased translational fidelity requires decreased translation rate, i.e. the “speed-accuracy trade-off” (26, 27). This model derives primarily from studies of codon•anticodon proofreading in *E. coli* and has been challenged in some studies (28, 29). To test if this applies to the discrimination of mischarged tRNAs by **9787**, we measured protein synthesis rates in *M. smegmatis* cultures. We compared untreated cultures to those treated with **9787** or with chloramphenicol, a known translation inhibitor. As expected, chloramphenicol dramatically reduced the rate of protein synthesis (**Fig. 8**). In contrast, cultures treated with **9787** at a concentration known to enhance fidelity showed translation rates that were not significantly different from untreated controls (**Fig. 8**). This key result demonstrates that **9787** enhances translational fidelity through a mechanism that is independent of the overall translation rate, challenging the universality of the speed-accuracy trade-off.

**Figure 7.**
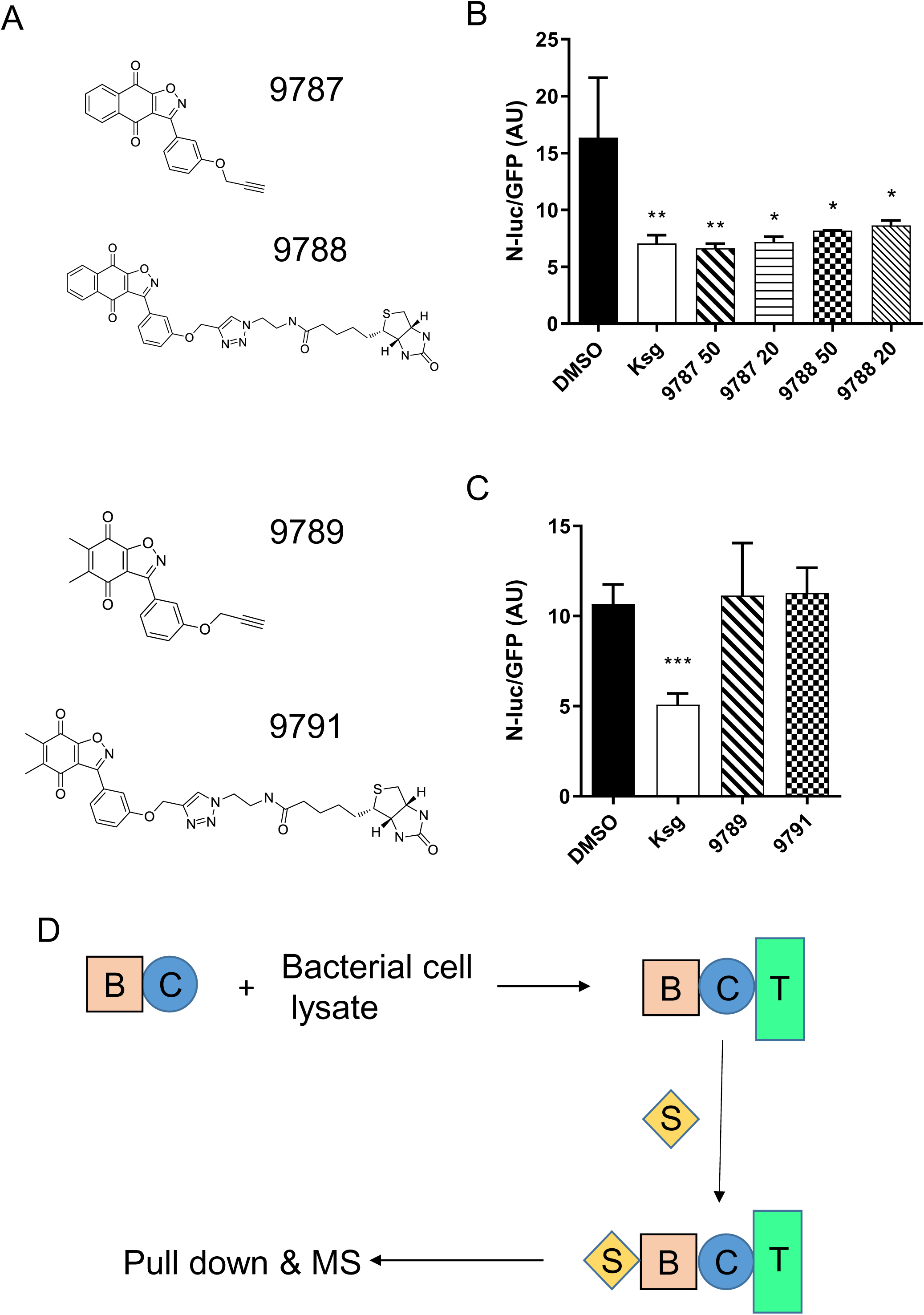
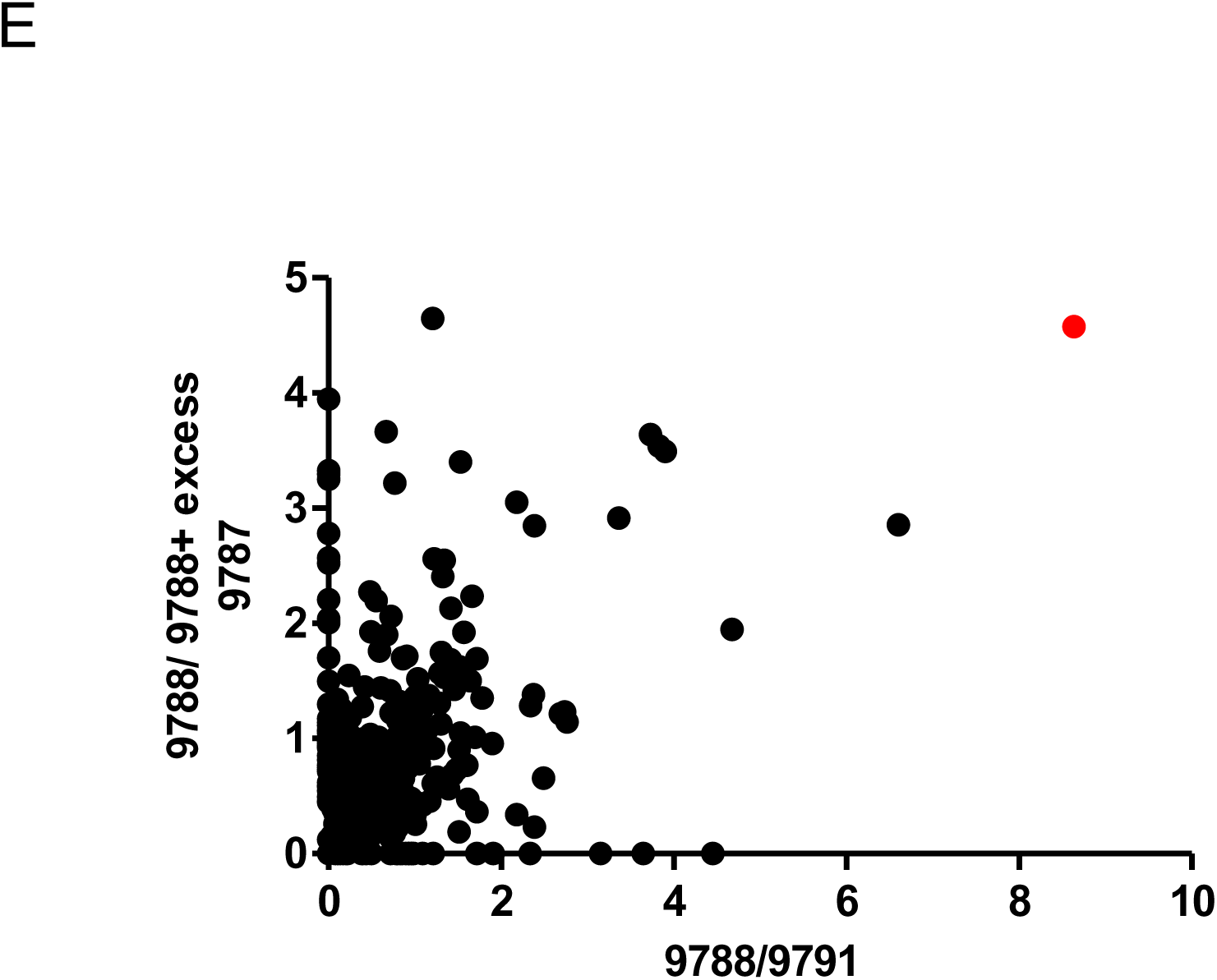
30S ribosomal protein S5 (RpS5) is the molecular target of compound 9787. (A) Structures of 9787, active biotinylated analog 9787, inactive analog 9789, and inactive biotinylated analog 9791. (B) The N-luc/GFP reporter was used to show the ability of 9787 and biotinylated 9788 to inhibit mycobacterial mistranslation. *p < 0.05, **p < 0.01, ***p < 0.001 by Student’s t-test (C) 9789, unable to reduce mycobacterial mistranslation, was biotinylated to produce compound 9791. Neither of the compounds could inhibit mycobacterial mistranslation. (D) Schematic of the experiment for target identification of 9787. The compound of interest (C) is tagged with biotin (B) and incubated with mycobacterial cell lysate. After it binds with the target (T), the whole complex is pulled down using streptavidin (S) beads (Materials & Methods for details). The pulled-down proteins are then analyzed by mass-spectrometry. (E) The ratio of the quality scores of the proteins pulled down by the active biotinylated compound (9788) and the inactive biotinylated compound (9791) has been plotted against the ratio of the quality scores of proteins pulled down by the active alone (9788), and the mixture of 9788 and excess non-biotinylated active compound (9787). The 30S ribosomal protein S5 (Rps5) is represented by the red dot.

**Figure 8.**
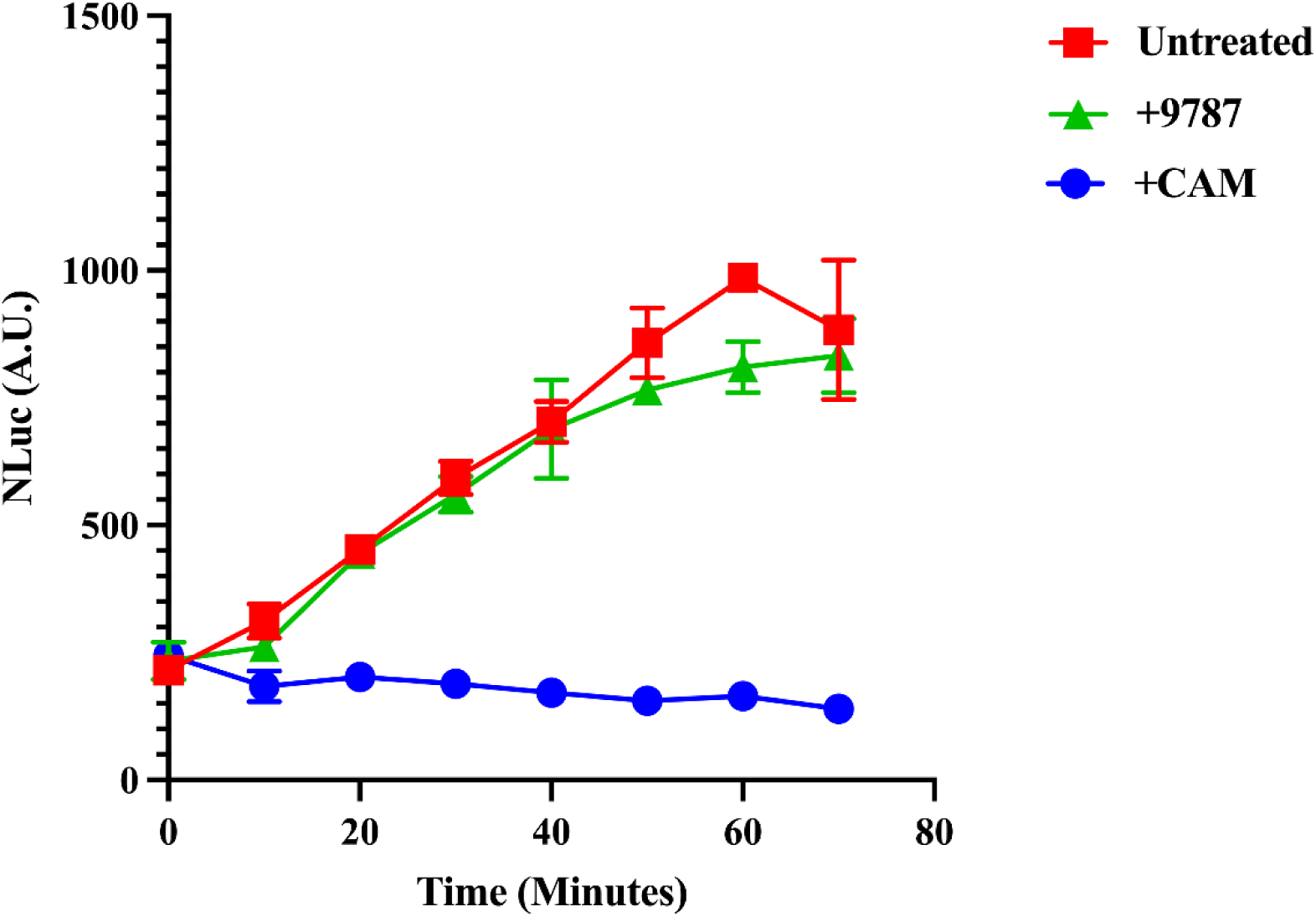
Treatment with compound 9787 does not alter the translation rate of *M. smegmatis*. Nanoluciferase (Nluc) activity was monitored over time as a proxy for translation rate in untreated *M. smegmatis,* and *M. smegmatis* treated with **9787** (100 µM) or chloramphenicol (200 µg/mL) using a strain of *M. smegmatis* in which Nluc expression was controlled by a tetracycline-inducible promoter. Time =0 represents time of addition of Atc (50ng/mL). Experiment was performed using three independent biological replicates. Points represent means +/-SD.

## DISCUSSION

In this study, we identify a novel class of synthetic molecules, benzo[d]isoxazole-4,7-diones, that increase mycobacterial translational fidelity. Our work makes three significant contributions. First, we establish a new chemical scaffold for modulating a specific mistranslation pathway that contributes to antibiotic tolerance. Second, using an unbiased chemical proteomics approach, we identify the 30S ribosomal protein S5 (RpS5) as the molecular target of our lead compound, **9787**. Third, we demonstrate that this compound can enhance translational fidelity without a concomitant decrease in the rate of protein synthesis, providing a clear exception to the canonical "speed-accuracy trade-off" model of translation.

The "speed-accuracy trade-off" has been considered a fundamental constraint on ribosomal function (26, 27, 30). This model proposes that ribosomes sacrifice speed to achieve high fidelity through kinetic proofreading of codon-anticodon interactions. Our finding that **9787** enhances fidelity without a measurable impact on translation rate suggests this trade-off is not absolute. This has several important implications. It suggests that the quality control mechanisms governing discrimination of mischarged tRNAs may be mechanistically distinct from those governing codon•anticodon matching. Furthermore, it reveals an unexpected plasticity in ribosome function, where fidelity can be modulated without the obligate fitness cost of slowed protein synthesis. Third, from a potential therapeutic perspective, it shows that translational fidelity can be pharmacologically modulated without imposing the fitness cost of slowed protein synthesis.

Our chemical proteomic approach identified RpS5 as the molecular target of **9787**. RpS5 plays crucial roles in mRNA binding to the ribosome and proper decoding (31). Our results suggest that small molecule binding to RpS5 can enhance discrimination of mischarged aminoacyl-tRNAs. How might the small ribosomal subunit, primarily concerned with decoding, contribute to aminoacyl identity discrimination? The mechanism remains unknown, but we speculate that binding of **9787** to RpS5 induces a conformational change that allosterically tightens quality control at the A-site, enhancing rejection of near-cognate or misacylated tRNAs without affecting the rate-limiting steps of translation. Several lines of evidence support a role for the small subunit in this process. First, kasugamycin, our previously identified mistranslation inhibitor (19), also targets the 30S subunit, binding near residues A794 and G926 of 16S rRNA (32, 33). The universally conserved nature of these residues suggests kasugamycin likely binds similarly in mycobacteria. Thus, the 30S ribosomal subunit appears to be a common target for mistranslation inhibitors. Second, while the ribosome’s proofreading of codon•anticodon interactions is well-established (34), its ability to detect mischarged tRNAs is less understood. Our results suggest the 30S subunit can discriminate aminoacyl identity, possibly through conformational sensing mechanisms that are independent of the kinetic proofreading governing codon•anticodon fidelity. Third, the lack of impact of **9787** on translation rate suggests that it acts independently of peptide bond formation or aa-tRNA accommodation, a rate limiting step in protein synthesis. Future studies, including structural or kinetic approaches will be crucial in understanding this mechanism.

Although maintaining translational fidelity is believed to be essential for all living systems, bacteria have been shown to tolerate up to 10% mistranslation of a specific codon without major deleterious effects (35). Moreover, mistranslation has been proposed to be a mechanism by which bacteria such as *Mycobacterium* adapt to environmental stresses such as low pH and presence of rifampicin. Under such conditions, mistranslation results in increased phenotypic heterogeneity and improves bacterial survival (29, 36).

Rifampicin is a crucial antibiotic in anti-TB chemotherapy. Its introduction in TB therapy has drastically reduced the treatment time. However, due to persistence of the bacteria, the current treatment time for drug-susceptible TB is still 4-6 months, and thus has room for further improvement. Empirical attempts have been made improve the efficacy of rifampicin by changing the therapeutic concentrations, as well as by using new combinations with other drugs (37-39). We, for the first time, had earlier proposed a mechanism of enhancing the efficacy of rifampicin in TB treatment by targeting mycobacterial mistranslation (19). In this work we have further expanded our search for compounds that can decrease mycobacterial mistranslation. The ‘hits’ have the potential to be used as candidates for combination therapy with rifampicin in future. Importantly, this gives us confidence of using our platform to develop future drug candidates. Moreover, the 30S ribosomal subunit can be used as an attractive drug target for reducing the burden of antibiotic-tolerant bacteria.

Phenotypic screening has been extensively used in developing novel drug candidates against *Mycobacterium tuberculosis* (40-43). Phenotypic screening is in general considered a better approach than biochemical screening, which often yields ‘hits’ that cannot actually penetrate into the bacterial cell. This is particularly true for mycobacteria that have a complex capsule and cell wall. On the other hand, hits obtained from whole cell phenotypic screening exhibit their effect after penetrating into the bacterial cell. Hence, they are usually better candidates for drug development than those obtained from biochemical screening.

In this study we have successfully expanded one of the luciferase-based mycobacterial mistranslation assays (19) developed in our laboratory into a larger scale screening platform. Throughout the screening procedure, kasugamycin was used as a positive control to enhance the validity of the screen. Although the performance of the screening platform was variable due to low average rate of mistranslation (∼1%/Asn codon), it nonetheless identified several hits that could be further validated and characterized. Most of the hits also significantly decreased mycobacterial RSPR. This further confirms the connection between mistranslation and RSPR in mycobacteria.

Several important questions remain: (1) Do these compounds affect other types of translational errors beyond mischarged tRNAs? (2) What is the structural basis for RpS5-mediated fidelity enhancement? (3) Can these compounds or optimized derivatives show efficacy in animal models of tuberculosis?

In conclusion, we have identified synthetic small molecules that increase mycobacterial translational fidelity through a rate-independent mechanism by targeting ribosomal protein S5. This work challenges prevailing models of ribosomal quality control, reveals new therapeutic strategies for antibiotic-tolerant bacteria, and demonstrates that the 30S ribosomal subunit is an attractive drug target for reducing the burden of persistent mycobacterial infections.

## MATERIALS & METHODS

### Bacterial strains, culture, antibiotics and test compounds

Wildtype *M. smegmatis* mc^2^-155 and its mistranslation reporter derivative strain were grown in Middlebrook 7H9 broth supplemented with 0.2% glycerol, 0.05% Tween-80, 10% Albumin-dextrose-salt (ADS), and on Luria-Bertani agar (LB agar) for plate assays. Rifampicin and kasugamycin (Ksg) were purchased from Sigma. Rifampicin was dissolved in dimethyl sulphoxide (DMSO); Ksg was dissolved in water, and filter sterilized. The test compounds for screening were all dissolved in DMSO.

### Screening of compounds that decrease mycobacterial mistranslation

Screening of the chemical library to identify compounds that decrease mycobacterial mistranslation was done using the previously developed N-luc/GFP mistranslation assay.^11^ Briefly, the mistranslation reporter *M. smegmatis* strain or *M. bovis* BCG strain was grown in 7H9 broth containing hygromycin (50µg/mL) and kanamycin (20µg/mL) at 37°C to late log phase (OD_600_ = 2.0). N-luc expression was induced by adding anhydrotetracycline (ATc, 50 ng/mL), and the bacterial culture was immediately aliquoted into a 96 well plate (100 µl, i.e. approximately 6x10^7^ cells per well). 50 µM of each of the test compound or 50 µg/mL of Ksg as positive control were added, and the cultures incubated for 16 hours at 37°C with shaking. Then the cultures were transferred to a 96-well black plate and GFP fluorescence was measured by Fluoroskan Ascent FL Fluorometer and Luminometer (Thermo Scientific). The plate was then centrifuged (4000 rpm for 10 minutes), and supernatants transferred to a 96-well white luminescence plate. The nano-luciferase assay was performed using Nano-Glo luciferase assay kit (Promega), and luminescence measured by the same machine.

### Rifampicin specific phenotypic resistance (RSPR) assay

RSPR assay was performed as previously described (19). Briefly, *M. smegmatis* mc^2^-155 was grown to stationary phase, after which the cultures were serially diluted and spread on LB-agar plates containing rifampicin (50µg/mL) only, or rifampicin and 50µM of each test compound. LB agar containing rifampicin and Ksg (100µg/mL) was used as positive control. The plates were incubated at 37°C for 5 days after which the number of colony forming units (cfu) were enumerated. The number of bacterial cells in the inoculum was calculated by counting colonies after spreading serial dilutions of the culture on LB-agar plates without any antibiotic.

### Target identification of 9788

To identify the intracellular target(s) of compounds that decrease mycobacterial mistranslation, 9788, a biotinylated version of a compound 9787 that decreased mycobacterial mistranslation, and 9791, a biotinylated version of a compound 9789 that cannot decrease mycobacterial mistranslation were used. A protocol modified from an earlier study to identify mycobacterial drug targets was used (44). To prepare mycobacterial cell lysate, *M. smegmatis* mc^2^-155 was grown in 7H9 broth up to OD_600_ 1.0, and then centrifuged at 4000 rpm for 10 minutes. The pellet was resuspended in phosphate-buffered saline (PBS) containing a tablet of protease inhibitor (Roche), and the cells were lysed by beads beating twice for 45 seconds each. Then the mixture was centrifuged at 15000 rpm for 15 minutes, the supernatant was collected and added to EZ view^TM^ Red streptavidin affinity gel (Sigma Aldrich), pre-equilibrated with the lysis buffer (PBS plus protease inhibitor cocktail). The mixture was incubated at 4°C for an hour in an orbital shaker and then centrifuged at 15000 rpm for an hour to collect the supernatant. The supernatant was then distributed to five tubes, and the protein concentration of each tube was adjusted to 1 mg/mL. The following compounds were then added one to each tube: 1) 1% DMSO, 2) 100 μM 9788, 3) 100 μM 9788 + 500 μM 9787, 4) 100 μM 9791, 5) 100 μM 9791 + 500 μM 9789. After 1 hour incubation at room temperature, the mixture from each tube was added to pre-equilibrated EZ view^TM^ Red streptavidin affinity gel, and incubated at 4°C for an hour in an orbital shaker. Then the mixtures were centrifuged at 15000 rpm for 20 mins at 4°C. The pellets were then resuspended in Laemmli buffer, ran in SDS PAGE, and sent for mass-spectrometry at the Tsinghua University Mass-Spectrometry facility.

### Relative Translation Rate Assay

To assess the relative translation rate, we adapted a previously described protocol(45) with minor modifications. *M. smegmatis* strain harboring the reporter construct was first grown in 7H9 medium to a stationary phase (OD₆₀₀ > 3). Cultures were then diluted to an OD₆₀₀ of 0.2 in fresh 7H9 medium, and anhydrotetracycline (ATc) was added to a final concentration of 50 ng/mL to induce reporter expression. After 5 minutes of induction, cells were pelleted, washed twice with fresh 7H9 medium (without ATc), and resuspended in the same medium supplemented with or without 200 μg/mL chloramphenicol (a translation inhibitor) and 100 μM compound **9787**. Higher concentrations of **9787** could not be tested since the stock solution was 10mM dissolved in DMSO and final DMSO concentration needed to be ≤1%. Aliquots were collected every 10 minutes and immediately snap-frozen in liquid nitrogen. Once all time points were collected, luminescence was measured using a luminometer. All experiments were performed in triplicate using three independent biological replicates.

## Author Contributions

MM and QWK optimized and performed the initial screen. JW performed all medicinal chemistry. SC performed follow-up microbiological experiments including target identification. GL provided the initial screening library and supervised JW. HJ performed the translation rate assay. SCF re-analyzed and interpreted all primary data, wrote the first and revised drafts of the manuscript and compiled all the figures. BJ conceived of the project, supervised the research and interpreted results, obtained funding and revised the manuscript.

## Acknowledgements

We thank the Tsinghua University proteomics core for performing the mass-spec analysis. This work was in part funded by the Gates Foundation (OPP1109789), a Wellcome Trust Investigator Award (207487/C/17/Z), a Kleberg Medical Research Foundation award (A141660) and startup funds from UCSF to BJ.

---

**Supplementary Table 1. Pull-down mass spectrometry to identify the binding target of 9787.** Putative binding proteins identified by mass spectrometry after mycobacterial cell lysates were incubated with (**A)** DMSO, **(B)** 9791, inactive biotinylated compound, **(C)** 9791 + 9789, inactive biotinylated compound with excess inactive non-biotinylated compound, **(D)** 9788, active biotinylated compound, and **(E)** 9788 + 9787, active biotinylated compound with excess active non-biotinylated compound.

## Notes

### Competing Interest Statement

The authors have declared no competing interest.

### Summary of Updates

Additional results (new Fig. 8) and extensively rewritten manuscript. Funding information added.

