## Supplementary material for "A small molecule increases mycobacterial translation fidelity without a fitness cost by targeting ribosomal protein S5": Supp information

**Supporting Information: The experimental procedures of chemistry and spectroscopic data of all compounds.**

**Title:** Medicinal chemistry of benzo[d]isoxazole-4,7-dione analogues identifies the ribosomal small subunit as a target for specific mycobacterial translational fidelity

**Authors:** Jie Wu^1^, Swarnava Chaudhuri^2^, Sarah Caldwell Feid^3^, Miaomiao Pan^2^, Qingwen Kawaji^2^, Gang Liu^1*^ and Babak Javid^2, 3*^

JW, SC and SCF contributed equally to this work

GL and BJ contributed equally to this work

**Running title**: Benzo[d]isoxazole-4,7-dione analogues decrease mycobacterial mistranslation

**MATERIALS**

Unless otherwise noted, commercially available materials were used without further purification. All NMR experiments were carried out on a Bruker Advance 400 MHz spectrometer using DMSO-*d_6_*, methanol-*d_4_*, Acetone-*d_6_* and CDCl_3_ as the solvent. Chemical shifts are given in ppm (*δ*) relative to the residual solvent and coupling constants (*J*) in Hz. Multiplicity is tabulated as s for singlet, d for doublet, t for triplet, q for quadruplet, and m for multiplicity. Melting points were determined without correction with a Yanaco micromelting point apparatus. UPLC-MS analysis was performed on a Thermo Finnigan LCQ Advantage mass spectrometer equipped with an Agilent pump, an Agilent detector, an Agilent liquid handler, and a fluent splitter. The column is a Kromasil C18 column (4.6 μm, 4.6 mm × 50 mm). The eluent was a mixture of acetonitrile and water containing 0.05% HCOOH with a linear gradient from 5:95 (v/v) to 95:5 (v/v) of acetonitrile-H_2_O within 5 minutes at a 0.3 mL/min flow rate for analysis. The UV detection was carried out at a UV wavelength of 254 nm. The 5% of the eluent was split into the MS system. Mass spectra were recorded in either positive or negative ion mode using electrospray ionization (ESI). High resolution LC-MS (HRMS) was carried out by Agilent LC/ MSD TOF using a column of Agilent ZORBAX SB-C18 (rapid resolution, 3.5 μm, 2.1 mm × 30 mm) at a flow of 0.40 mL/min. The solvent was methanol/water = 75/25 (v/v) containing 5 mmol/L ammonium formate. The ion source was electrospray ionization (ESI). The column chromatography was performed with silica gel 60 (200-300 mesh) from Qingdao Haiyang Chemical Factory. The tested compounds were purified ≥ 95%, as characterized by HPLC under UV 254 nm wavelength, NMR and HRMS.

**EXPERIMENTAL PROCEDURES OF CHEMISTRY AND SPECTROSCOPIC DATA**

**General Procedure one:** To a solution of aldehyde (10.0 mmol) in methanol (50 mL) was added potassium carbonate (11.0 mmol) and hydroxylamino hydrochloride (11.0 mmol) at room temperature. The mixture was stirred at room temperature for 3 h until the reaction was complete. The solution was evaporated in vacuo and the residue was stirred in ethyl acetate (50.0 mL) for 10 minutes. The solution was filtered and the filtrate was evaporated to get the intermediate aldehyde oxime.

To a solution of phenol (2.0 mmol) in acetonitrile (10 mL) and water (5 mL) was added PhI(OAc)_2_ (4.0 mmol) at room temperature. The mixture was stirred at room temperature until the starting material disappeared. Then aldehyde oxime (3.0 mmol) was added. After the flask was completely cooled in an ice bath, 4.0 mmol of PhI(OAc)_2_ solved in acetonitrile (8 mL) and water (2mL) was added dropwise within 0.5 h and then the mixture warmed to room temperature. After the intermediate disappeared, the mixture was poured into brine (60 mL) and extracted with ethyl acetate (50 mL × 3). The combined organic layer was washed with brine (50 mL × 2) and dried over Na_2_SO_4_. After evaporation of the solvent in vacuo, the compound was obtained and characterized after purification by silica gel column chromatography (ethyl acetate/petroleum ether).

**5,6-dimethyl-3-phenylbenzo[d]isoxazole-4,7-dione (7006)**. Yellow powder in 80.1% yield. ^1^H-NMR (400 MHz, CDCl_3_) *δ* 8.11 (d, *J* = 7.5 Hz, 2H), 7.49-7.56 (m, 3H), 2.17 (s, 6H). ^13^C-NMR (100 MHz, CDCl_3_) *δ*180.7, 175.3, 164.9, 160.2, 144.1, 140.1, 131.2, 129.2, 128.6, 126.2, 117.5, 13.10, 12.04. ESI-MS: calcd for C_15_H_11_NO_3_ [M+H]^+^: 254.12; found, 254.07.

**3-(4-chlorophenyl)-5,6-dimethylbenzo[d]isoxazole-4,7-dione (7013)**. Yellow powder in 85.0% yield. ^1^H-NMR (400 MHz, CDCl_3_) *δ* 8.12 (d, *J* = 8.4 Hz, 2H), 7.49 (d, *J* = 8.4 Hz, 2H), 2.17 (s, 6H). ^13^C-NMR (100 MHz, CDCl_3_) *δ* 180.7, 175.1, 165.0, 159.2, 144.1, 140.3, 137.5, 130.5, 129.0, 124.7, 117.4, 13.1, 12.1. ESI-MS: calcd for C_15_H_10_ClNO_3_ [M+H]^+^: 288.19; found, 288.03.

**3-(4-fluorophenyl)-5,6-dimethylbenzo[d]isoxazole-4,7-dione (7020)**. Yellow powder in 69.8% yield. ^1^H-NMR (400 MHz, CDCl_3_) *δ* 8.18 (dd, *J* = 8.6 Hz, 5.5 Hz, 2H), 7.20 (t, *J* = 8.6 Hz, 2H), 2.17 (s, 6H). ^13^C-NMR (100 MHz, CDCl_3_) *δ* 180.8, 175.2, 165.8, 165.0, 163.3, 159.2, 144.1, 140.3, 131.5, 131.4, 122.4, 122.4, 117.3, 116.0, 115.8, 13.1, 12.1. ESI-MS: calcd for C_15_H_10_FNO_3_ [M+H]^+^: 272.54; found, 272.06.

**5,6-dimethyl-3-(naphthalen-1-yl)benzo[d]isoxazole-4,7-dione (9719)**. Yellow powder in 61.4% yield. ^1^H-NMR (400 MHz, CDCl_3_) *δ* 8.04 (d, *J* = 8.2 Hz, 1H), 7.94 (d, *J* = 8.1 Hz, 1H), 7.86 (d, *J* = 8.3 Hz, 1H), 7.73 (d, *J* = 7.1 Hz, 1H), 7.61 – 7.49 (m, 3H), 2.17 (s, 3H), 2.07 (s, 3H). ^13^C-NMR (100 MHz, CDCl_3_) *δ* 180.3, 175.4, 164.3, 159.4, 143.8, 140.5, 133.6, 131.3, 131.1, 129.1, 128.6, 127.2, 126.5, 125.1, 124.9, 123.3, 118.9, 12.9, 12.2. ESI-MS: calcd for C_19_H_13_NO_3_ [M+H]^+^: 304.51; found, 304.09.

**3-(4-fluorophenyl)-5,6-dimethyl-1H-indazole-4,7-dione (9741)**. Yellow powder in 64.1% yield. ^1^H-NMR (400 MHz, DMSO-*d_6_*) *δ* 14.55 (s, 1H), 8.15 (s, 2H), 7.37 (s, 2H), 2.03 (s, 6H). ^13^C-NMR (100 MHz, DMSO-*d_6_*) *δ* 181.3, 179.6, 164.6, 162.1, 144.8, 140.9, 130.7, 130.6, 125.2, 115.4, 115.3, 115.1, 13.1, 12.2. ESI-MS: calcd for C_15_H_11_FN_2_O_2_ [M+H]^+^: 271.37; found, 271.08.

**5,6-dichloro-3-(p-tolyl)benzo[d]isoxazole-4,7-dione (9744)**. Yellow powder in 53.1% yield. ^1^H-NMR (400 MHz, CDCl_3_) *δ* 7.95 (d, *J* = 8.2 Hz, 1H), 7.74 (d, *J* = 8.2 Hz, 1H), 7.31 (dd, *J* = 8.0 Hz, 2.9 Hz, 2H), 2.44 (d, *J* = 2.7 Hz, 3H). ^13^C-NMR (100 MHz, CDCl_3_) *δ* 170.3, 169.5, 159.9, 159.3, 143.5, 142.0, 129.8, 129.4, 129.2, 127.3, 117.4, 21.6. ESI-MS: calcd for C_14_H_7_Cl_2_NO_3_ [M+H]^+^: 308.36; found, 307.98.

**3-(o-tolyl)naphtho[2,3-d]isoxazole-4,9-dione (9759).** Yellow powder in 83.4% yield. ^1^H-NMR (400 MHz, CDCl_3_) *δ* 8.36-8.28 (m, 1H), 8.24-8.19 (m, 1H), 7.90-7.83 (m, 2H), 7.48 (dd, *J* = 12.2 Hz, 4.5 Hz, 2H), 7.44-7.33 (m, 2H), 2.37 (s, 3H). ^13^C-NMR (100 MHz, CDCl_3_) *δ* 178.5, 173.4, 165.4, 160.8, 137.6, 135.2, 134.3, 133.5, 132.1, 130.66, 130.65, 130.2, 127.49, 127.43, 125.7, 125.5, 120.6, 20.2. ESI-MS: calcd for C_18_H_11_NO_3_ [M+H]^+^: 290.26 found, 290.07.

**3-(naphthalen-1-yl)naphtho[2,3-d]isoxazole-4,9-dione (9761)**. Yellow powder in 93.1% yield. ^1^H-NMR (400 MHz, CDCl_3_) *δ* 8.36-8.30 (m, 1H), 8.15 (dt, *J* = 6.6 Hz, 3.1 Hz, 1H), 8.08 (d, *J* = 8.3 Hz, 1H), 7.97 (d, *J* = 7.7 Hz, 1H), 7.89 (d, *J* = 8.4 Hz, 1H), 7.87-7.76 (m, 3H), 7.63 (dd, *J* = 8.2 Hz, 7.2 Hz, 1H), 7.51-7.57 (m, 2H). ^13^C-NMR (100 MHz, CDCl_3_) *δ* 178.2, 173.4, 165.6, 160.2, 135.3, 134.3, 133.6, 133.5, 132.1, 131.4, 131.2, 129.1, 128.6, 127.5, 127.4, 127.2, 126.5, 125.0, 124.9, 123.2, 121.0. ESI-MS: calcd for C_21_H_11_NO_3_ [M+H]^+^: 326.58 found, 326.07.

**3-cyclohexylnaphtho[2,3-d]isoxazole-4,9-dione (9767)**. Yellow powder in 89.3% yield. ^1^H-NMR (400 MHz, CDCl_3_) *δ* 8.29-8.19 (m, 2H), 7.87-7.79 (m, 2H), 3.26-3.30 (m, 1H), 2.09 (d, *J* = 11.4 Hz, 2H), 1.93-1.86 (m, 2H), 1.81-1.67 (m, 3H), 1.40-1.46 (m, 3H). ^13^C-NMR (100 MHz, CDCl_3_) *δ* 179.4, 173.4, 166.3, 165.3, 135.1, 134.2, 133.5, 132.1, 127.4, 127.3, 120.1, 36.1, 30.5, 26.0, 25.7. ESI-MS: calcd for C_17_H_15_NO_3_ [M+H]^+^: 282.34 found, 282.11.

**3-(2,4-dimethylphenyl)naphtho[2,3-d]isoxazole-4,9-dione (9775)**. Yellow powder in 81.2% yield. ^1^H-NMR (400 MHz, CDCl_3_) *δ* 8.29 (dd, *J* = 6.0 Hz, 2.9 Hz, 1H), 8.22-8.15 (m, 1H), 7.87-7.81 (m, 2H), 7.38 (d, *J* = 7.8 Hz, 1H), 7.21-7.11 (m, 2H), 2.41 (s, 3H), 2.32 (s, 3H). ^13^C-NMR (100 MHz, CDCl_3_) *δ* 178.5, 173.4, 165.4, 160.8, 140.7, 137.4, 135.2, 134.2, 133.6, 132.1, 131.5, 130.2, 127.4, 127.3, 126.5, 122.5, 120.6, 21.4, 20.1. ESI-MS: calcd for C_19_H_13_NO_3_ [M+H]^+^: 304.54 found, 304.09.

**3-(3-bromo-4-methylphenyl)naphtho[2,3-d]isoxazole-4,9-dione (9782)**. Yellow powder in 73.1% yield. ^1^H-NMR (400 MHz, CDCl_3_) *δ* 8.42 (s, 1H), 8.29 (dd, *J* = 7.4 Hz, 1.4 Hz, 2H), 8.06 (dd, *J* = 7.9 Hz, 1.4 Hz, 1H), 7.90-7.82 (m, 2H), 7.41 (d, *J* = 7.9 Hz, 1H), 2.50 (s, 3H). ^13^C-NMR (100 MHz, CDCl_3_) *δ* 178.5, 173.2, 166.3, 159.6, 141.5, 135.3, 134.3, 133.7, 132.9, 131.7, 130.9, 128.1, 127.8, 127.3, 125.4, 125.1, 119.5, 23.1. ESI-MS: calcd for C_18_H_10_BrNO_3_ [M+H]^+^: 368.02 found, 367.98.

**3-(2-methoxy-3-methylphenyl)naphtho[2,3-d]isoxazole-4,9-dione (9783)**. Yellow powder in 43.6% yield. ^1^H-NMR (400 MHz, CDCl_3_) *δ* 8.30-8.10 (m, 2H), 7.90-7.72 (m, 2H), 7.46-7.32 (m, 2H), 7.32-7.16 (m, 1H), 3.99 (s, 3H), 2.66 (s, 3H). ^13^C-NMR (100 MHz, CDCl_3_) *δ* 183.0, 180.7, 162.4, 159.3, 151.4, 134.6, 133.7, 131.7, 131.2, 130.4, 126.7, 124.3, 122.2, 120.9, 119.3, 117.1, 61.8, 15.2. ESI-MS: calcd for C_19_H_13_NO_4_ [M+H]^+^: 320.18 found, 320.08.

**3-(3-(prop-2-yn-1-yloxy)phenyl)naphtho[2,3-d]isoxazole-4,9-dione (9787)**. Yellow powder in 80.1% yield. ^1^H-NMR (400 MHz, DMSO-*d_6_*) *δ* 8.20 (d, *J* = 7.0 Hz, 2H), 7.99 (dt, *J* = 6.9 Hz, 5.7 Hz, 2H), 7.73 (d, *J* = 7.1 Hz, 2H), 7.57 (t, *J* = 8.0 Hz, 1H), 7.31 – 7.25 (m, 1H), 4.93 (d, *J* = 2.2 Hz, 2H), 3.64 (t, *J* = 2.1 Hz, 1H).^13^C-NMR (100 MHz, CDCl_3_) *δ* 178.5, 173.3, 166.3, 160.7, 157.6, 135.3, 134.3, 133.8, 131.7, 129.8, 127.8, 127.4, 127.2, 122.6, 119.6, 118.4, 115.5, 78.2, 75.8, 56.1. ESI-MS: calcd for C_20_H_11_NO_4_ [M+H]^+^: 330.24 found, 330.07.

**5,6-dimethyl-3-(3-(prop-2-yn-1-yloxy)phenyl)benzo[d]isoxazole-4,7-dione (9789)**. Yellow powder in 81.1% yield. ^1^H-NMR (400 MHz, CDCl_3_) *δ* 7.62-7.36 (m, 2H), 7.29 (t, *J* = 3.0 Hz, 1H), 7.00-7.12 (m, 1H), 4.97 (d, *J* = 2.1 Hz, 2H), 2.61 (s, 1H), 2.12 (s, 3H), 2.17 (s, 3H). ^13^C-NMR (100 MHz, CDCl_3_) *δ* 178.5, 173.3, 166.3, 160.7, 157.6, 135.3, 134.3, 133.8, 131.7, 129.8, 127.8, 127.4, 127.2, 122.6, 119.6, 118.4, 115.5, 78.2, 75.8, 56.1. ^13^C-NMR (100 MHz, CDCl_3_) *δ* 183.7, 175.9, 161.5, 160.4, 156.0, 144.9, 143.6, 131.0, 128.7, 122.8, 116.5, 113.8, 112.6, 79.6, 76.8, 58.0, 12.7, 12.2. ESI-MS: calcd for C_18_H_13_NO_4_ [M+H]^+^: 308.53 found, 308.08.

**General Procedure two:** To a solution of benzo[d]isoxazole-4,7-dione (1.2 mmol) in dichloromethane (15.0 mL) was added Na_2_O_4_S_2_ (7.1 mmol) in water (15.0 mL) at room temperature. The mixture was stirred at room temperature until the starting material disappeared. Then the mixture was poured into brine (60 mL) and extracted with dichloromethane (20 mL × 3). The combined organic layer was washed with brine (50 mL × 2) and dried over Na_2_SO_4_. After evaporation of the solvent in vacuo, the intermediate was dissolved in acetone (15 mL). Then K_2_CO_3_ (4.1 mmol) and CH_3_I or CH_3_CH_3_I (4.7 mmol) were added. The mixture was stirred at room temperature for 3 h until the reaction was complete. The solution was filtered and the filtrate was evaporated to get the crude product. The compound was obtained and characterized after purification by silica gel column chromatography (ethyl acetate/petroleum ether).

**4,7-dimethoxy-5,6-dimethyl-3-phenylbenzo[d]isoxazole (9735)**. White solid in 78.9% yield. ^1^H-NMR (400 MHz, DMSO-*d_6_*) *δ* 7.88 (dd, *J* = 7.1, 2.0 Hz, 2H), 7.63-7.56 (m, 3H), 4.03 (s, 3H), 3.24 (s, 3H), 2.29 (s, 3H), 2.22 (s, 3H). ^13^C-NMR (100 MHz, DMSO-*d_6_*) *δ* 157.5, 155.4, 146.4, 137.9, 132.2, 130.6, 129.4, 128.9, 128.5, 126.3, 113.8, 62.1, 60.9, 13.2, 12.5. ESI-MS: calcd for C_17_H_17_NO_3_ [M+H]^+^: 254.62; found, 284.12.

**3-(4-chlorophenyl)-4,7-dimethoxy-5,6-dimethylbenzo[d]isoxazole (9736)**. White solid in 84.6% yield. ^1^H-NMR (400 MHz, DMSO-*d_6_*) *δ* 7.93 (d, *J* = 8.4 Hz, 2H), 7.66 (d, *J* = 8.4 Hz, 2H), 4.03 (s, 3H), 3.29 (s, 3H), 2.29 (s, 3H), 2.23 (s, 3H). ^13^C-NMR (100 MHz, DMSO-*d_6_*) *δ* 156.5, 155.4, 146.3, 137.9, 135.5, 132.4, 131.3, 129.1, 127.3, 126.5, 113.6, 62.2, 60.9, 13.2, 12.5. ESI-MS: calcd for C_17_H_16_ClNO_3_ [M+H]^+^: 318.52; found, 318.08.

**3-(4-fluorophenyl)-4,7-dimethoxy-5,6-dimethylbenzo[d]isoxazole (9737)**. White solid in 84.9% yield. ^1^H-NMR (400 MHz, DMSO-*d_6_*) *δ* 7.95 (dd, *J* = 8.4 Hz, 5.6 Hz, 2H), 7.43 (t, *J* = 8.8 Hz, 2H), 4.03 (s, 3H), 3.27 (s, 3H), 2.29 (s, 3H), 2.22 (s, 3H). ^13^C-NMR (100 MHz, DMSO-*d_6_*) *δ* 164.9, 162.5, 156.6, 155.4, 146.3, 137.9, 132.3, 131.9, 131.8, 126.3, 124.9, 124.9, 116.2, 115.9, 113.7, 62.1, 60.5, 13.2, 12.5. ESI-MS: calcd for C_17_H_16_FNO_3_ [M+H]^+^: 302.18; found, 302.11.

**4,7-diethoxy-3-(4-fluorophenyl)-5,6-dimethylbenzo[d]isoxazole (9738)**. White solid in 80.2% yield. ^1^H-NMR (400 MHz, DMSO-*d_6_*) *δ* 8.01-7.94 (m, 2H), 7.52-7.46 (m, 2H), 4.32 (q, *J* = 7.0 Hz, 2H), 3.46 (q, *J* = 7.0 Hz, 2H), 2.35 (s, 3H), 2.27 (s, 3H), 1.43 (t, *J* = 7.0 Hz, 3H), 1.00 (t, *J* = 7.0 Hz, 3H). ^13^C-NMR (100 MHz, DMSO-*d_6_*) *δ* 164.9, 162.5, 156.8, 155.6, 145.2, 136.7, 132.8, 132.1, 131.9, 126.6, 125.02, 124.99, 116.1, 115.9, 113.9, 70.9, 69.1, 15.9, 14.9, 13.6, 12.9. ESI-MS: calcd for C_19_H_20_FNO_3_ [M+H]^+^: 330.47; found, 330.14.

**4,7-dimethoxy-5,6-dimethyl-3-(p-tolyl)benzo[d]isoxazole (9739)**. White solid in 71.2% yield. ^1^H-NMR (400 MHz, DMSO-*d_6_*) *δ* 7.79 (d, *J* = 8.1 Hz, 2H), 7.39 (d, *J* = 7.9 Hz, 2H), 4.02 (s, 3H), 3.26 (s, 3H), 2.42 (s, 3H), 2.28 (s, 3H), 2.22 (s, 3H). ^13^C-NMR (100 MHz, DMSO-*d_6_*) *δ* 157.4, 155.4, 146.4, 140.3, 137.9, 132.1, 129.5, 129.3, 126.2, 125.6, 113.8, 62.2, 60.9, 21.4, 13.2, 12.5. ESI-MS: calcd for C_18_H_19_NO_3_ [M+H]^+^: 298.36; found, 298.14.

**4,9-dimethoxy-3-(p-tolyl)-5,6,7,8-tetrahydronaphtho[2,3-d]isoxazole (9746)**. White solid in 73.2% yield. ^1^H-NMR (400 MHz, DMSO-*d_6_*) *δ* 7.78 (d, *J* = 8.0 Hz, 2H), 7.39 (d, *J* = 7.9 Hz, 2H), 4.05 (s, 3H), 3.26 (s, 3H), 2.85- 2.68 (m, 4H), 2.42 (s, 3H), 1.82-1.65 (m, 4H). ^13^C-NMR (100 MHz, DMSO-*d_6_*) *δ* 157.3, 154.4, 146.0, 140.3, 137.2, 132.2, 129.6, 129.2, 126.7, 125.6, 113.7, 61.8, 60.4, 24.1, 23.3, 22.1, 22.1, 21.4. ESI-MS: calcd for C_20_H_21_NO_3_ [M+H]^+^: 324.47; found, 324.25.

**5,6-dichloro-4,7-dimethoxy-3-(p-tolyl)benzo[d]isoxazole (9747)**. White solid in 8.7% yield. ^1^H-NMR (400 MHz, Acetone-*d_6_*) *δ* 7.83 (d, *J* = 8.2 Hz, 2H), 7.44 (d, *J* = 7.9 Hz, 2H), 4.26 (s, 3H), 3.51 (s, 3H), 2.46 (s, 3H). ^13^C-NMR (100 MHz, Acetone-*d_6_*) *δ*158.5, 156.1, 146.4, 141.6, 138.7, 130.1, 126.7, 125.5, 122.8, 116.9, 62.6, 61.6, 21.4. ESI-MS: calcd for C_16_H_13_Cl_2_NO_3_ [M+H]^+^: 338.07; found, 338.03.

**3-(4-fluorophenyl)benzo[d]isoxazole (9748)**. To a solution of 4-fluoro-N-hydroxybenzimidoyl chloride (1.2 mmol), anthranilic acid (2.3 mmol) and K_2_CO_3_ (1.7 mmol) in CH_3_CN (15.0 mL) was added tert-butyl nitrite (2.3 mmol) at room temperature. The mixture was stirred at 115^o^C for 20 h. Then the mixture was evaporated to get the crude product. The compound was obtained and characterized after purification by silica gel column chromatography (ethyl acetate/petroleum ether). Yellow powder in 41.3% yield. ^1^H-NMR (400 MHz, CDCl_3_) *δ* 8.03-7.92 (m, 2H), 7.90 (d, *J* = 8.0 Hz, 1H), 7.60-7.64 (m, 2H), 7.43-7.37 (m, 1H), 7.24-7.29 (m，2H). ^13^C-NMR (100 MHz, CDCl_3_) *δ* 165.2, 163.9, 162.7, 156.4, 130.0, 129.9, 129.8, 125.15, 125.12, 124.0, 121.9, 120.3, 116.4, 116.2, 110.2. ESI-MS: calcd for C_13_H_8_FNO [M+H]^+^: 214.51; found, 214.06.

**N-(2-(4-((3-(4,9-dioxo-4,9-dihydronaphtho[2,3-d]isoxazol-3-yl)phenoxy)methyl)-1H-1,2,3-triazol-1-yl)ethyl)-5-((3aS,4S,6aR)-2-oxohexahydro-1H-thieno[3,4-d]imidazol-4-yl)penta-namide (9788)**. To a solution of **9787** (1.0 mmol) and N-(2-azidoethyl)-5-((3aS,4S,6aR)-2-oxohexahydro-1H-thieno[3,4-d]imidazol-4-yl)pentanamide (1.2 mmol) in N,N-dimethylfo-rmamide (15.0 mL) and water (10.0 mL) was added CuSO_4_ (1.5 mmol) and sodium ascorbate (0.5 mmol) at room temperature. The mixture was stirred at 43^o^C for 10 h. Then the mixture was poured into brine (60 mL) and extracted with ethyl acetate (20 mL × 3). The combined organic layer was washed with brine (50 mL × 2) and dried over Na_2_SO_4_. After evaporation of the solvent in vacuo, the compound was obtained and characterized after purification by silica gel column chromatography (ethyl acetate/petroleum ether). Yellow powder in 53.5% yield. ^1^H-NMR (400 MHz, DMSO-*d_6_*) *δ* 8.25-8.27 (m, 3H), 8.01 (s, 3H), 7.79-7.82 (m, 1H), 7.68 (d, *J* = 6.3 Hz, 1H), 7.56 (s, 1H), 7.33-7.36 (m, 1H), 6.40 (d, *J* = 27.3 Hz, 1H), 5.26 (s, 1H), 4.45 (s, 2H), 4.20-4.23 (m, 2H), 3.51 (s, 2H), 3.36 (s, 3H), 3.08 (s, 1H), 2.79 (s, 1H), 2.03 (d, *J* = 6.3 Hz, 2H), 1.56 (s, 1H), 1.45-1.47 (m, 3H), 1.23-1.26 (m, 3H). ^13^C-NMR (100 MHz, DMSO-*d_6_*) *δ* 179.1, 173.7, 173.0, 167.1, 165.1, 160.5, 158.5, 143.2, 135.7, 134.9, 134.1, 132.3, 130.5, 127.7, 127.6, 127.5, 127.0, 122.0, 119.4, 117.9, 116.1, 61.8, 61.4, 59.6, 55.8, 49.4, 47.3, 35.4, 28.5, 28.4, 25.6. ESI-MS: calcd for C_32_H_31_N_7_O_6_S [M+H]^+^: 642.46 found, 642.21.

**N-(2-(4-((3-(5,6-dimethyl-4,7-dioxo-4,7-dihydrobenzo[d]isoxazol-3-yl)phenoxy)methyl)-1H-1,2,3-triazol-1-yl)ethyl)-5-((3aS,4S,6aR)-2-oxohexahydro-1H-thieno[3,4-d]imidazol-4-yl)pe-ntanamide (9791)**. Brown powder in 30.5% yield. ^1^H-NMR (400 MHz, DMSO-*d_6_*) *δ* 7.50-7.52 (m, 1H), 7.42 (t, *J* = 14.8 Hz, 1H), 7.27 (t, *J* = 3.0 Hz, 1H), 7.15 (s, 1H), 6.98-7.11 (m, 1H), 6.43 (s, 1H), 5.61 (t, *J* = 10.2 Hz, 2H), 5.55 (s, 1H), 5.42 (s, 1H), 5.16 (s, 2H), 4.57-4.61 (m, 1H), 4.48 (dd, *J* = 18.5, 16.1 Hz, 1H), 3.71 (t, *J* = 10.2 Hz, 2H), 3.45-3.33 (m, 3H), 2.35 (dt, *J* = 15.2, 11.6 Hz, 1H), 2.25-2.18 (m, 1H), 2.17-2.07 (m, 2H), 1.87 (s, 6H), 1.78-1.62 (m, 2H), 1.32-1.16 (m, 2H). ^13^C-NMR (100 MHz, DMSO-*d_6_*) *δ* 184.5, 176.7, 174.8, 163.7, 161.8, 160.5, 159.2, 143.0, 142.2, 141.7, 131.2, 128.8, 122.8, 121.9, 116.6, 112.6, 109.0, 68.1, 60.9, 56.6, 55.2, 49.9, 38.0, 37.4, 36.9, 29.9, 26.6, 25.3, 12.7. ESI-MS: calcd for C_30_H_33_N_7_O_6_S [M+H]^+^: 622.71 found, 620.52.
